# MTCH2 drives metabolic remodeling in muscle satellite cells via mitochondrial fatty acid utilization

**DOI:** 10.64898/2026.11.17.766893

**Authors:** Akira Murakami, Yuta Aoyagi, Ayane Miyazawa, Takashi Masuda, Miki Suzuki, Kohjiro Nagao, Masaki Tsuchiya, Yusuke Ono, Kotaro Hirano, Masato Umeda, Yuji Hara

## Abstract

Muscle satellite cell (MuSC) differentiation into myoblasts underpins skeletal muscle regeneration by supporting myofiber repair and formation. This process involves extensive metabolic remodeling from glycolysis to mitochondrial oxidative metabolism. Although this switch is required for proper MuSC differentiation, its molecular regulators remain unclear. Here, we identify mitochondrial carrier homolog 2 (MTCH2) as a regulator of this remodeling. MTCH2 contributes preferentially to MuSC differentiation rather than proliferation and supports mitochondrial respiration, particularly under a low-glucose condition. Loss of Mtch2 fragments mitochondria and suppresses respiration associated with fatty acid β-oxidation. Notably, MTCH2-dependent fatty acid utilization appears largely independent of carnitine palmitoyltransferase 1 (CPT1). Furthermore, skeletal muscle regeneration is impaired in MuSC-specific *Mtch2* conditional knockout mice. Together, these findings establish MTCH2 as a regulator of mitochondrial metabolic remodeling in MuSCs and highlight its context-dependent roles in cellular metabolism and function.

**Summary Statement:** Mitochondrial MTCH2 regulates metabolic remodeling during muscle satellite cell differentiation. Our findings identify a role for MTCH2 in skeletal muscle regeneration and highlight its context-dependent functions.

## Introduction

Skeletal muscle homeostasis relies on the capacity to regenerate myofibers (Wagers and Conboy, 2005). Myofiber regeneration depends on the myogenic function of muscle satellite cells (MuSCs): myoblasts that differentiate from MuSCs fuse to form multinucleated myotubes (Relaix and Zammit, 2012). Understanding the molecular mechanisms that govern myogenesis is therefore important to both cell biology and regenerative biology.

During myogenic differentiation, cells undergo extensive metabolic remodeling, including a shift from glycolysis to mitochondrial oxidative metabolism (Relaix et al., 2021). The energetic demands of differentiation require increased mitochondrial activity, including fatty acid β-oxidation (Folmes et al., 2012). *In vitro*, low-serum culture conditions trigger this metabolic switch and promote myotube formation (Brennan and Olson, 1990; Rochard et al., 2000). Despite widespread recognition of the importance of metabolic reprogramming during myogenesis, the molecular mechanisms coordinating mitochondrial metabolism, particularly fatty acid β-oxidation, remain poorly understood. These observations point to the existence of molecular regulators that drive the metabolic switch accompanying myogenic differentiation.

Mitochondrial carrier homolog 2 (MTCH2) belongs to the solute carrier (SLC) 25 family and is located in the outer mitochondrial membrane, although its transporter function and substrates remain unclear (Ruprecht and Kunji, 2020). Studies have linked MTCH2 to diverse cellular processes, including apoptosis (Grinberg et al., 2005; Zaltsman et al., 2010), and mitochondrial metabolism and dynamics (Bahat et al., 2018; Goldman et al., 2024; Labbé et al., 2021; Maryanovich et al., 2015). Its involvement in mitochondrial function and cell fate regulation suggests that MTCH2 may link metabolic state to differentiation programs. However, its contribution to myogenesis and the accompanying metabolic remodeling has yet to be explored.

In this study, we investigated the role of MTCH2 in MuSC differentiation using *in vitro* and *ex vivo* systems and assessed its physiological significance *in vivo*. Our findings show that MTCH2 promotes myogenesis and muscle regeneration by supporting mitochondrial function in myoblasts. Our analyses further revealed that MTCH2 supports mitochondrial respiration through the promotion of fatty acid β-oxidation.

## Results

### MTCH2 is required for myogenic differentiation

Analysis of published transcriptomic datasets from a mouse muscle injury model (Harada et al., 2018) revealed that *Mtch2* mRNA expression increases during MuSC activation and myogenic differentiation (Supplementary Fig. 1A). We also observed a progressive increase in *Mtch2* expression in C2C12 myoblasts cultured under low-serum conditions (Supplementary Fig. 1B). These findings prompted us to investigate the role of MTCH2 in myogenesis.

We then assessed how siRNA-mediated MTCH2 depletion affects myogenic differentiation in C2C12 cells. *Mtch2* knockdown significantly reduced the expression of the myogenic differentiation marker *Myog*, the myotube maturation markers *Myh4* and *Mck* (Arnold and Winter, 1998) and the fusion factor *Mymk* (Millay et al., 2013) (Fig. 1A). Accordingly, *Mtch2*-deficient cells showed impaired myotube formation, reflected in a reduced fusion index (Fig. 1B). By contrast, *Mtch2* knockdown did not significantly affect cell proliferation under growth conditions (Fig. 1C). We observed similar effects in primary myoblasts derived from MuSCs isolated from mice: MTCH2 depletion impaired myotube formation without affecting proliferation (Fig. 1D, E). Independent siRNAs targeting *Mtch2* yielded similar results (Supplementary Fig. 2). Conversely, *Mtch2* overexpression enhanced myotube formation under low-serum conditions (Supplementary Fig. 3). Together, these results demonstrate that MTCH2 is required for myogenic differentiation.

**Figure 1.**
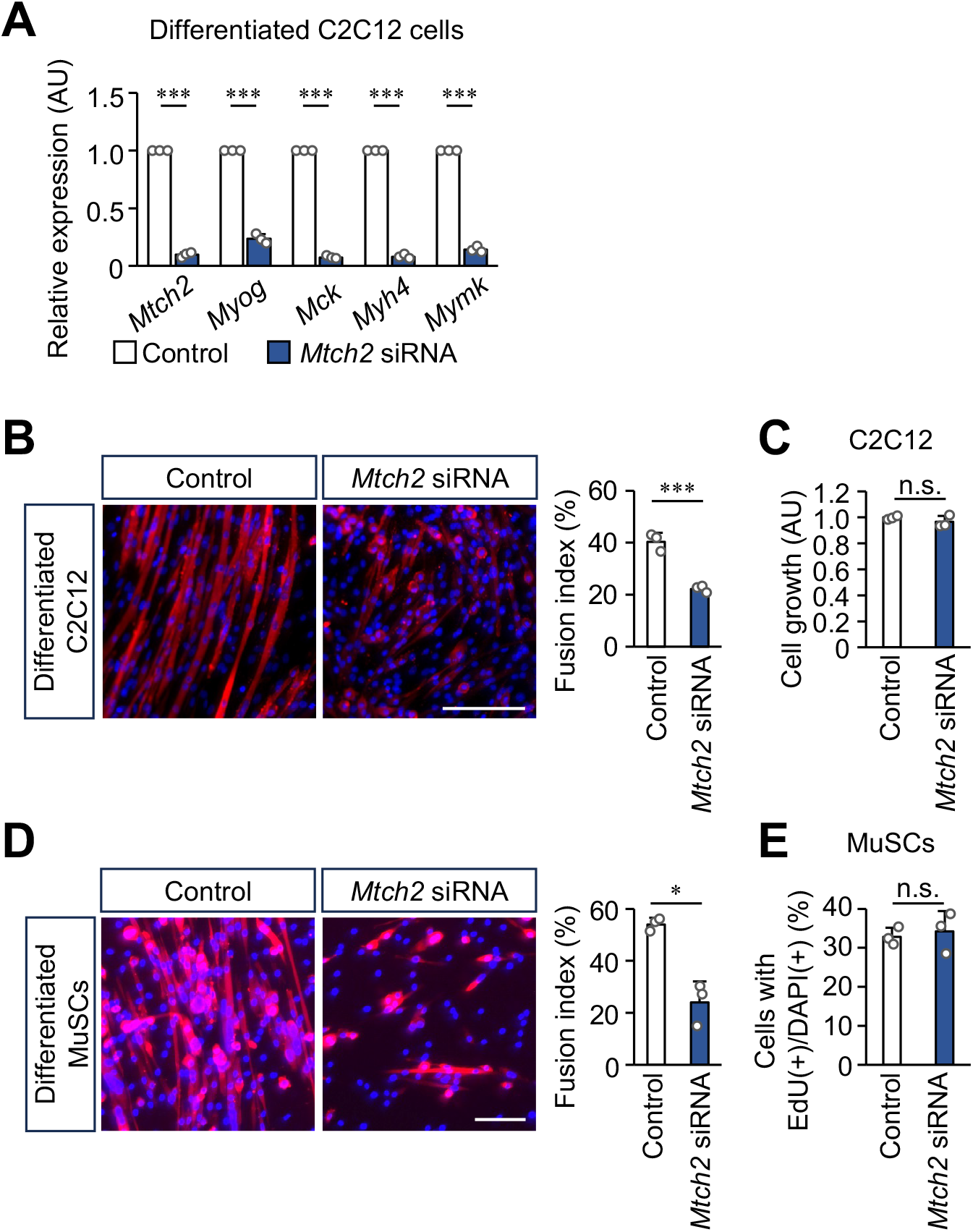
Myogenic differentiation in *Mtch2*-depleted myoblasts. (A) Expression of myogenic differentiation marker mRNAs in C2C12 cells after 2 days of culture in low-serum medium. *Myog*, myogenin; *Mck*, muscle creatine kinase; *Myh4*, myosin heavy chain 4; *Mymk*, myomaker. (B) Myotube formation in C2C12 cells cultured for 4 days in low-serum medium, quantified using the fusion index. (C) C2C12 cell growth assessed with the 3-(4,5-Dimethyl-2-thiazolyl)-2,5-diphenyltetrazolium Bromide (MTT) assay. (D) Myotube formation in myoblasts derived from isolated MuSCs after 2 days of culture in low-serum medium, quantified using the fusion index. (E) Growth of myoblasts derived from isolated MuSCs, assessed by EdU incorporation 6 days after isolation. *Mtch2* depletion was achieved using siRNA. Red, myosin heavy chain; blue, nuclei. Scale bars, 100 μm. Data are presented as mean + SD. n = 3. \**P* < 0.05, \*\**P* < 0.01, \*\*\**P* < 0.001; n.s., not significant.

### MTCH2 is involved in mitochondrial function during myogenesis

To investigate the molecular basis of MTCH2 function in myogenesis, we used RNA-seq to analyze primary myoblasts derived from isolated MuSCs after siRNA-mediated MTCH2 depletion under low-serum conditions. Gene set enrichment analysis (GSEA) showed significant downregulation of genes involved in mitochondrial function, particularly mitochondrial respiration, in *Mtch2*-depleted cells compared with controls (Fig. 2A). We validated these findings by quantitative RT-PCR, confirming reduced expression of representative genes in the affected pathways (Fig. 2B). Together, these findings suggest that MTCH2 is required to support mitochondrial function during myogenesis.

**Figure 2.**
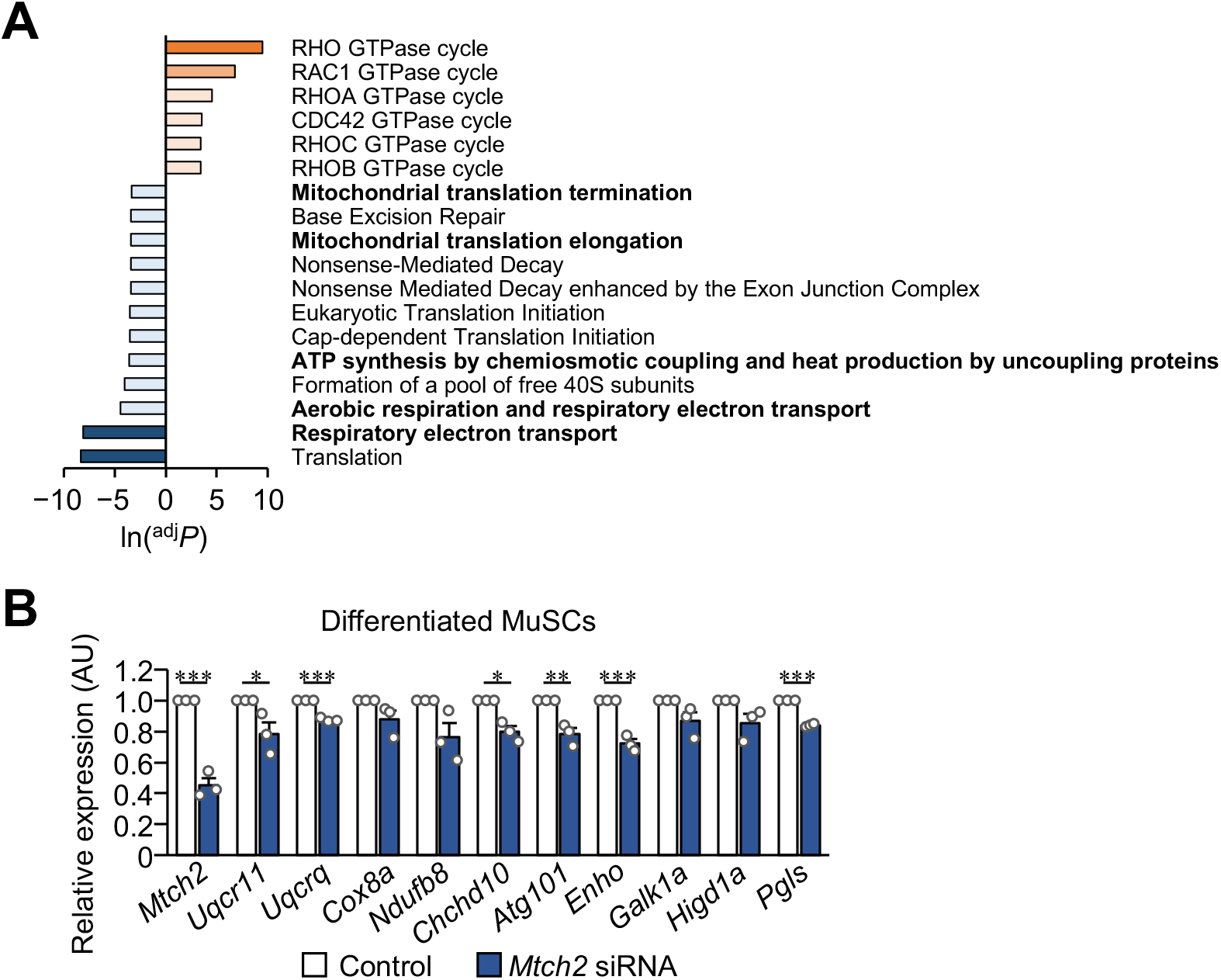
Transcriptomic profile of *Mtch2*-depleted primary myoblasts. (A) Gene set enrichment analysis of myoblasts differentiated from isolated MuSCs following *Mtch2* depletion. (B) Quantitative real-time PCR validation of the genes identified by GSEA as downregulated in *Mtch2*-depleted cells in (A). Cells were cultured for 2 days in low-serum medium before RNA-seq or quantitative real-time PCR analysis. *Mtch2* depletion was achieved using siRNA. Data are presented as mean + SD. n = 3. \**P* < 0.05, \*\**P* < 0.01, \*\*\**P* < 0.001.

### MTCH2 promotes mitochondrial respiration under low-glucose conditions

We next investigated how MTCH2 contributes to mitochondrial function in myoblasts. Under standard culture conditions, MTCH2 suppression increased mitochondrial fragmentation relative to control cells (Fig. 3A). Despite this morphological change, however, MTCH2 suppression did not significantly alter the oxygen consumption rate (OCR) under these conditions (Fig. 3B). To determine whether a respiratory defect emerges when reliance on glycolysis is reduced, we cultured cells in low-glucose medium. Under these conditions, both basal OCR and maximal OCR measured after treatment with carbonyl cyanide 4-(trifluoromethoxy)phenylhydrazone (FCCP) were significantly lower in *Mtch2*-depleted cells than in controls (Fig. 3C). These findings indicate that MTCH2 promotes mitochondrial respiration in myoblasts, with its effect becoming apparent when glucose availability is limited.

**Figure 3.**
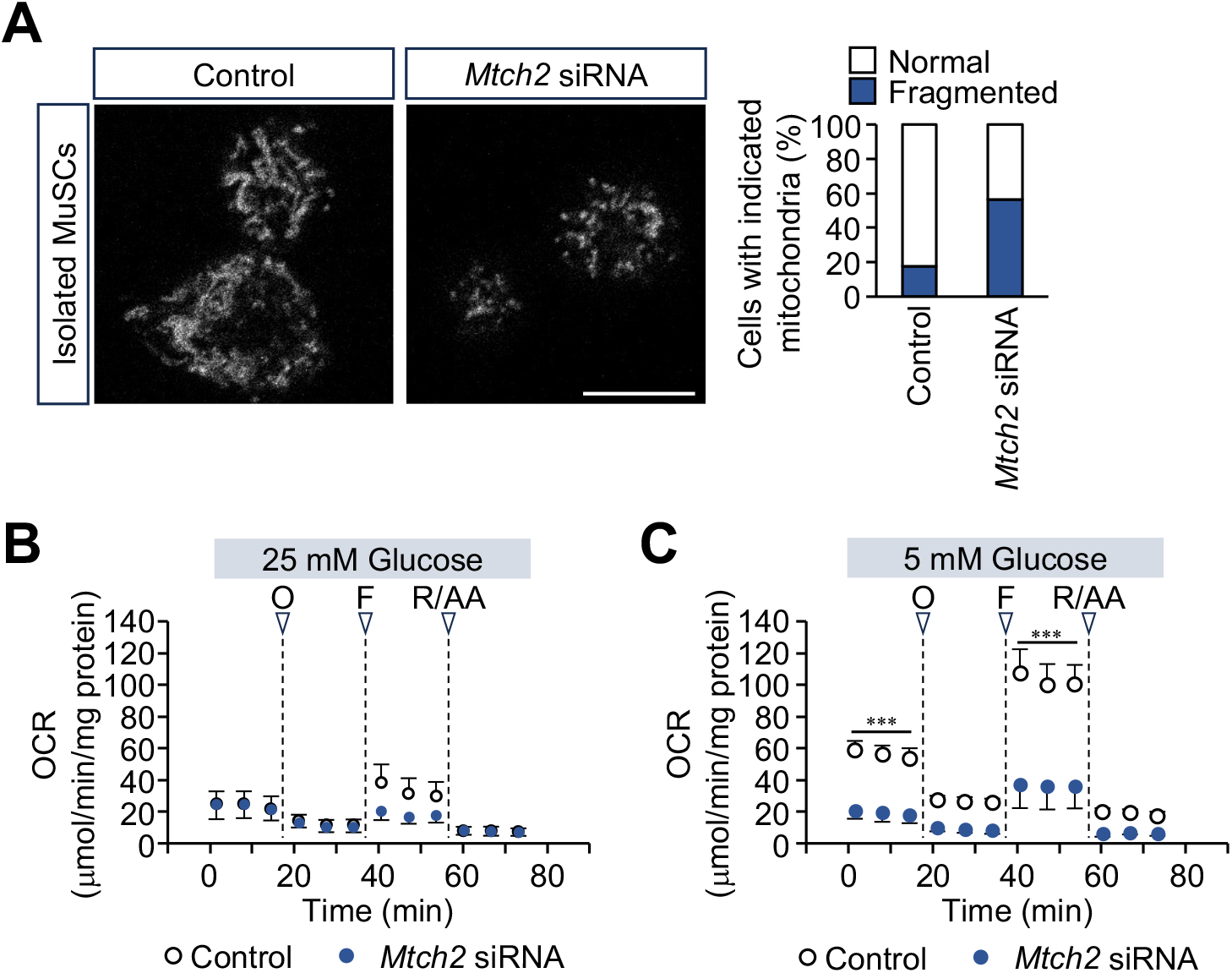
Mitochondrial morphology and respiration in *Mtch2*-depleted primary myoblasts. (A) Morphology of mitochondria in myoblasts differentiated from isolated MuSCs following *Mtch2* depletion. Mitochondria were stained with MitoTracker® and classified as fused/elongated (Normal) or fragmented (Fragmented). Control, n = 216; *Mtch2* siRNA, n = 229; N = 3 mice each. Scale bar, 10 μm. (B, C) Oxygen consumption rate (OCR) in *Mtch2*-depleted MuSC-derived myoblasts cultured in assay medium with 25 mM glucose (B) or 5 mM glucose (C). O, oligomycin; F, FCCP (carbonyl cyanide 4-(trifluoromethoxy)phenylhydrazone); R/AA, rotenone/antimycin A. Cells were subjected to analysis on day 6 after isolation. *Mtch2* depletion was achieved using siRNA. Data are presented as mean + SD. N = 3 mice. \*\*\**P* < 0.001.

### MTCH2 is involved in CPT1-independent fatty acid β-oxidation in myoblasts

To determine how MTCH2 regulates mitochondrial respiration, we assessed the relative contributions of glycolysis and fatty acid oxidation. We measured the extracellular acidification rate (ECAR) in C2C12 cells under a series ofmetabolic stress conditions (Wu et al., 2007). Basal and maximal glycolytic activities were comparable in control and *Mtch2*-depleted cells (Fig. 4A). We then assessed fatty acid oxidation by measuring OCR in cells cultured in low-glucose medium supplemented with long-chain fatty acids. OCR was significantly lower in *Mtch2*-depleted cells than in controls under these conditions (Fig. 4B). Notably, this difference persisted in the presence of etomoxir, an inhibitor of carnitine palmitoyltransferase 1 (CPT1), a rate-limiting enzyme in fatty acid β-oxidation that mediates fatty acid transport into mitochondria (Lopaschuk et al., 1988), whereas etomoxir-sensitive OCR was comparable between groups (Fig. 4C). These findings indicate that MTCH2 promotes mitochondrial respiration primarily through fatty acid oxidation rather than glycolysis. They further suggest that MTCH2 regulates fatty acid utilization through a CPT1-independent mechanism.

**Figure 4.**
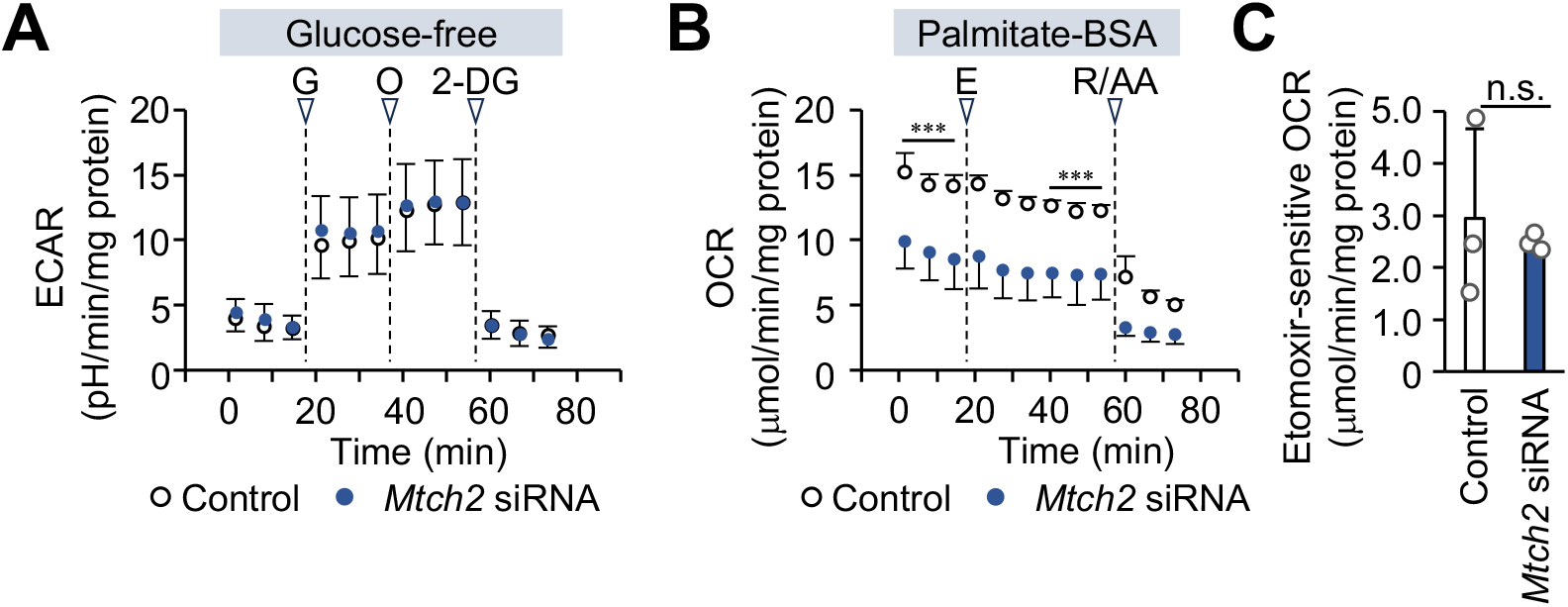
Glycolysis and fatty acid β-oxidation in *Mtch2*-depleted myoblasts. (A) Extracellular acidification rate (ECAR) in *Mtch2*-depleted C2C12 cells, measured in glucose-free assay medium. G, glucose; O, oligomycin; 2-DG, 2-deoxy-D-glucose. n = 3. (B) OCR dependent on fatty acid β-oxidation in *Mtch2*-depleted C2C12 cells. Measurements were performed in palmitate-containing assay medium following preincubation for 1 h in culture medium containing 0.5 mM glucose. n = 3. (C) Dependence of fatty acid β-oxidation-dependent OCR on carnitine palmitoyltransferase 1 (CPT1) in C2C12 cells. The etomoxir sensitivity of OCR was calculated from (B). Data are presented as mean + SD. n = 3. \*\*\**P* < 0.001; n.s., not significant.

### *In vivo Mtch2* loss in MuSCs impairs myogenic differentiation and alters mitochondrial morphology

To validate these findings with an independent genetic approach, we generated MuSC-specific *Mtch2* conditional knockout (cKO) mice (Supplementary Fig. 5A) and induced *Mtch2* deletion *in vivo* by administering tamoxifen intraperitoneally (Supplementary Fig. 5B). We subsequently isolated *Mtch2*-deficient MuSCs and differentiated them into myoblasts in culture for analysis. Quantitative real-time PCR confirmed efficient depletion of *Mtch2* expression (Supplementary Fig. 5C). *Mtch2*-deficient myoblasts had a significantly higher proportion of fragmented mitochondria than control cells (Fig. 5A). The percentage of MYOG-positive cells was also significantly lower in *Mtch2* cKO cells than in controls (Fig. 5B). Consistent with these findings, *Mtch2* cKO cells showed impaired myotube formation under low-serum differentiation conditions (Fig. 5C). Together, these results indicate that the defects following siRNA-mediated *Mtch2* depletion reflect the loss of MTCH2 function itself rather than siRNA-specific effects.

**Figure 5.**
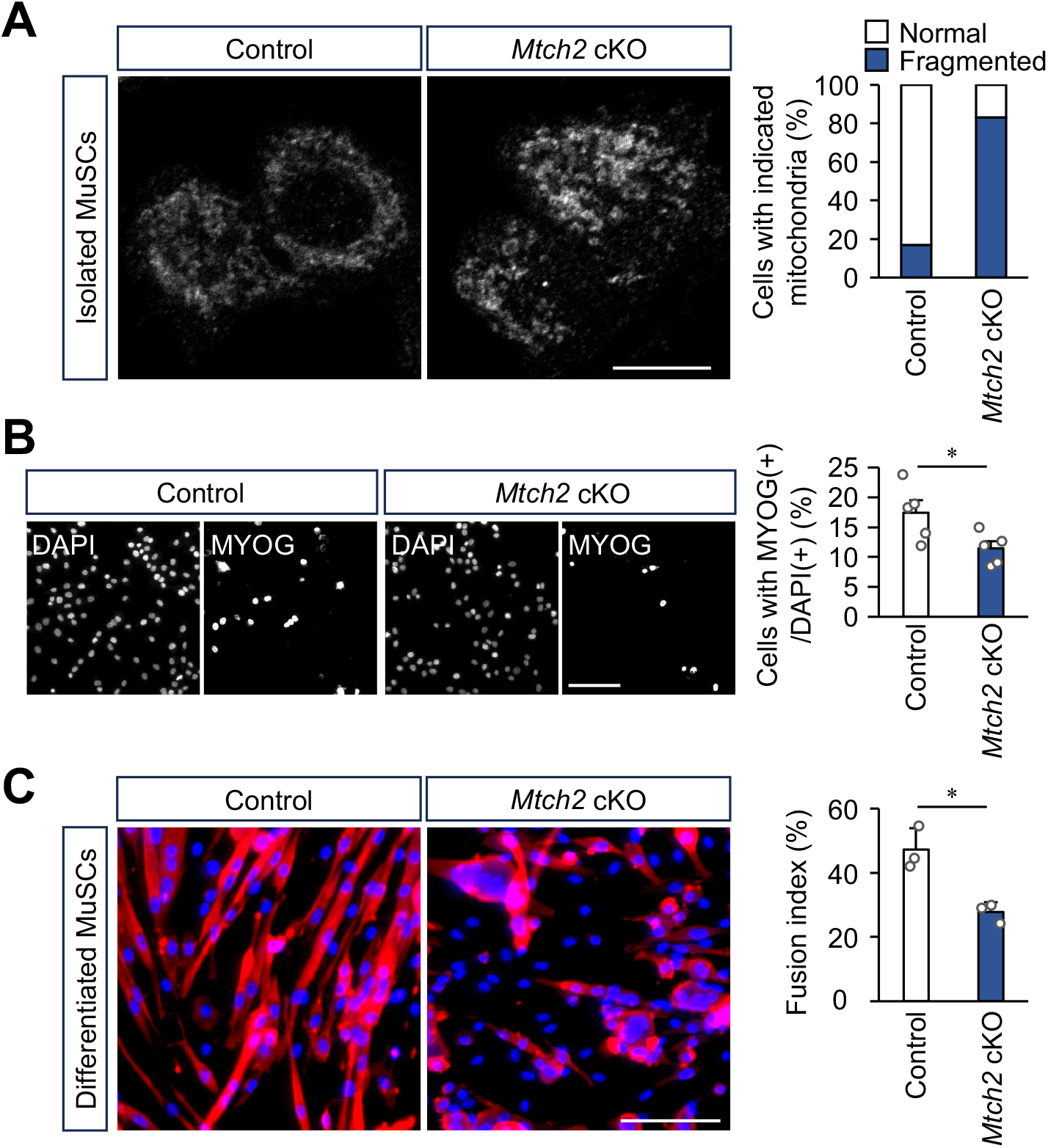
Characterization of primary myoblasts derived from MuSCs isolated from MuSC-specific *Mtch2* conditional knockout (*Mtch2* cKO) mice. (A) Mitochondrial morphology. Mitochondria were stained with MitoTracker® and classified as fused/elongated (Normal) or fragmented (Fragmented). Control, n = 391; *Mtch2* cKO, n = 438; N = 3 mice each. Scale bar, 10 μm. (B) Immunofluorescence analysis of MYOG expression. Scale bar, 100 μm. (C) Myotube formation in primary myoblasts after 2 days of culture in low-serum medium, quantified using the fusion index. Red, myosin heavy chain; blue, nuclei. Scale bars, 100 μm. Data are presented as mean + SD. N = 3 mice. MuSCs were isolated from control or *Mtch2* cKO mice, differentiated into myoblasts, and analyzed on day 5 after isolation. \**P* < 0.05.

### MTCH2 is required for muscle regeneration *in vivo*

Finally, we examined the role of MTCH2 in muscle regeneration in *Mtch2* cKO mice. After tamoxifen administration, we induced muscle injury by injecting cardiotoxin into the tibialis anterior (TA) muscle and collected tissues 7 days later (Fig. 6A). Histological analysis of TA muscles revealed that muscle regeneration was moderately affected in *Mtch2* cKO mice compared with control mice (Fig. 6B). Immunofluorescence staining for laminin further showed that regenerating myofibers had a smaller cross-sectional area in *Mtch2* cKO mice than in controls (Fig. 6C), indicating thinner myofibers. Together, these findings demonstrate that MTCH2 is required for muscle regeneration *in vivo* and support the physiological relevance of the *Mtch2*-dependent myogenesis observed ex vivo.

**Figure 6.**
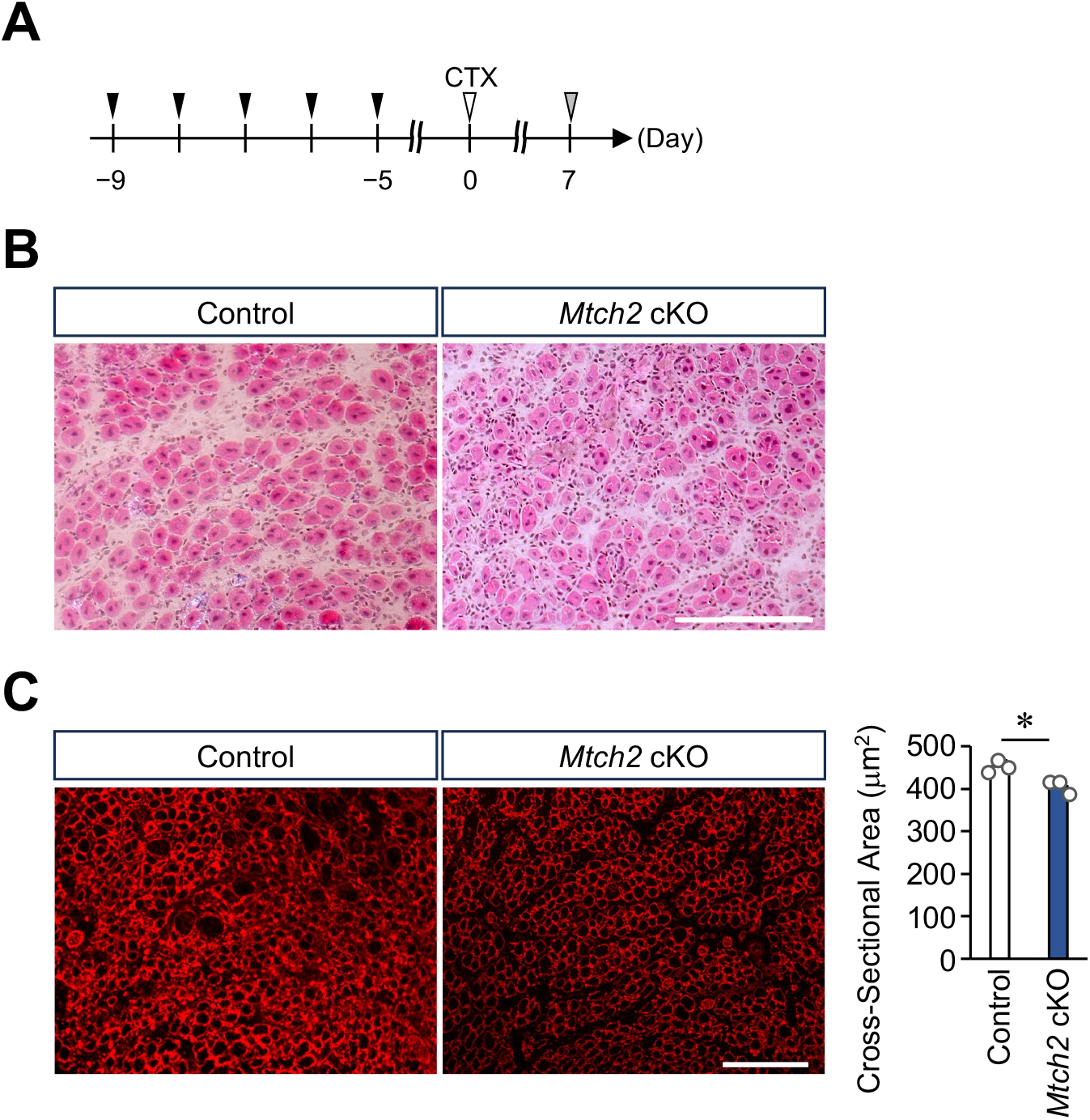
Skeletal muscle regeneration in *Mtch2* cKO mice. (A) Experimental scheme showing the induction of MuSC-specific *Mtch2* knockout and cardiotoxin (CTX)-induced muscle injury. Black arrows, intraperitoneal tamoxifen administration; gray arrows, skeletal muscle isolation. (B) Hematoxylin and eosin staining of cross-sections from CTX-injured TA muscles collected 7 days after injection. (C) Cross-sectional area of regenerating myofibers at 7 days after CTX injection. The left panel shows representative laminin-stained sections of regenerating muscle; the right panel quantifies the average myofiber cross-sectional area. Red, laminin. Scale bar, 100 μm. Data are presented as mean + SD. N = 3 mice. \**P* < 0.05.

## Discussion

A shift from glycolysis to mitochondrial β-oxidation is essential for energy production during myogenesis, but the molecular regulators coordinating this transition remain unclear. Although MTCH2 has been implicated in differentiation in several tissues, its contribution to MuSC differentiation remains unexplored. Here, *in silico* and *in vitro* analyses first indicated a potential role for MTCH2 in myogenesis, which we subsequently confirmed functionally in isolated MuSC-derived myoblasts. Our RNA-seq analysis suggests that MTCH2 contributes to myogenesis by regulating mitochondrial function. Indeed, MTCH2 supported mitochondrial respiration in MuSC-derived myoblasts, with a more pronounced contribution when glucose availability was limited. These observations led us to identify mitochondrial fatty acid β-oxidation as a process involving MTCH2. We confirmed these findings in MuSC-specific *Mtch2* cKO mice and cells isolated from them. Together, our results support a model in which MTCH2 mediates metabolic remodeling during myogenesis.

We found that loss of MTCH2 induced mitochondrial fragmentation in myoblasts. Consistent with this observation, previous studies have described fragmented, rounded mitochondria with reduced fusion activity (Bahat et al., 2018; Goldman et al., 2024). MTCH2 also cooperates with the mitochondrial fusion factor MFN2 to promote starvation-induced mitochondrial hyperfusion (Labbé et al., 2021). These findings suggest that MTCH2-dependent regulation of mitochondrial morphology may act upstream of myogenesis. The small-molecule MFN activator MASM7 was recently reported to enhance mitochondrial fusion (Zacharioudakis et al., 2022). In our experiments, MASM7 treatment suppressed mitochondrial fragmentation induced by *Mtch2* knockdown in myoblasts (Supplementary Fig. 6A). It also partially rescued the defect in myotube formation caused by MTCH2 loss (Supplementary Fig. 6B). These findings support a major role for the regulation of mitochondrial morphology in MTCH2-dependent myogenesis, although morphology-independent mechanisms are also likely to contribute.

We found that MTCH2 promotes mitochondrial fatty acid β-oxidation in myoblasts, particularly through a CPT1-independent mechanism. Previous studies have reported inconsistent effects of MTCH2 loss on mitochondrial respiration and metabolism (Bahat et al., 2018; Bar-Lev et al., 2016; Chourasia et al., 2025; Maryanovich et al., 2015; Ruggiero et al., 2017). Differences in how strongly the examined cell types rely on glycolysis versus β-oxidation to meet their energy demands may explain these discrepancies. Indeed, the effects of MTCH2 loss on mitochondrial respiration were more pronounced under a low-glucose condition (Fig. 3B, C). MTCH2 loss also preferentially impaired myotube formation under nutrient-poor, low-serum conditions rather than proliferation in nutrient-rich medium (Fig. 1B–D). Moreover, the low-glucose condition exacerbated the myogenic defects induced by MTCH2 loss (Supplementary Fig. 7). These observations show that nutrient availability and differentiation state influence how MTCH2 contributes to cellular function, even within a single cell type. Thus, the physiological role of MTCH2 and its relationship to CPT1 dependency may vary with cellular context, including cell type, metabolic state, and differentiation state (Wu et al., 2025). Our findings suggest that MTCH2 supports mitochondrial function during muscle regeneration *in vivo*; however, increased mitochondrial oxidative phosphorylation and mitochondrial content were reported in a study of myofiber-specific *Mtch2* knockout mice, seemingly in contrast to our observations (Buzaglo-Azriel et al., 2016). Differences in cellular state and experimental timescale may explain these contrasting findings. The previous study examined chronic *Mtch2* deficiency in mature myofibers, whereas we focused on relatively acute *Mtch2* depletion in MuSCs and differentiated myoblasts. Adaptive responses to chronic MTCH2 loss may also contribute to the phenotypes reported previously (Bahat et al., 2018). Together, these *in vivo* observations further suggest that MTCH2 function depends strongly on physiological context^1^. How the MTCH2-dependent enhancement of mitochondrial function observed *ex vivo* promotes muscle regeneration *in vivo* remains unclear. Experimental systems that allow precise control of metabolic and nutritional conditions *in vivo* will be needed to resolve this question.

Understanding the molecular mechanisms by which MTCH2 regulates myogenesis may offer new insights into skeletal muscle disorders. In particular, elucidating the mechanism of MTCH2-dependent, CPT1-independent fatty acid β-oxidation is an important direction for future research. One possibility is that MTCH2 directly mediates fatty acid transport across the mitochondrial membrane through a process analogous to CPT1; alternatively, it may exert broad, indirect control over unknown fatty acid transport pathways. The first possibility is consistent with a report suggesting that MTCH2 has phospholipid scramblase activity (Bartoš et al., 2024); the second is supported by a study proposing that MTCH2 acts as a mitochondrial outer membrane protein insertase (Guna et al., 2022). Through broader effects on mitochondrial quality control, including mitophagy and membrane protein homeostasis, MTCH2 might influence unidentified fatty acid oxidation pathways or transport systems. Consistent with this possibility, *Mtch2* knockdown reduced mitochondrial content in myoblasts (Supplementary Fig. 8). In addition to its association with obesity, MTCH2 has been reported to have important roles in multiple tissues (Escott-Price et al., 2014; Kulyté et al., 2011; Willer, 2009). Elucidating the molecular basis of MTCH2-mediated metabolic reprogramming may therefore advance both skeletal muscle biology and regenerative biology more broadly.

## Material and Methods

### Mice

Animal care, ethical use, and protocols were approved by the Animal Care Use and Review Committee of the University of Shizuoka. *Mtch2*^tm1a(EUCOMM)Hmgu^ mice were purchased from the European Conditional Mouse Mutagenesis Program (EUCOMM) (Skarnes et al., 2011). To remove the *lacZ* and neomycin resistance cassettes, *Mtch2*^tm1a^ mice were crossed with B6-Tg(CAG-FLPe)36 mice (RBRC01834; RIKEN BioResource Research Center [BRC]) (Kanki et al., 2006). The resulting *Mtch2*^tm1c^ mice were subsequently crossed with *Pax7 iRES-CreERT2/+* mice (The Jax laboratory; (Murphy et al., 2011)) to generate *Mtch2*^flox/flox^; *Pax7-IRES-CreERT2/+* mice. To induce Cre-mediated deletion of the floxed exon 6 and 7 in *Pax7*-expressing MuSCs, tamoxifen (TMX, #T5648, Sigma-Aldrich; 20 mg ml^−1^ in corn oil) was administered intraperitoneally at a dose of 40 μg g^−1^ body weight for five consecutive days.

### MuSC isolation using FACS

MuSCs were isolated from limb muscles as previously described (Hirano et al., 2025; Hirano et al., 2023; Murakami et al., 2026). Briefly, skeletal muscle samples from mouse forelimbs were digested with 0.2% collagenase type II (#CLS2; Worthington). Mononuclear cells were then incubated for at least 30 min at 4°C with PE-conjugated anti-mouse Ly-6A/E (Sca-1) antibody (#122508; BioLegend), PE-conjugated anti-mouse CD45 antibody (#103106; BioLegend), PE-conjugated anti-mouse CD31 antibody (#102508; BioLegend), and allophycocyanin-conjugated anti-mouse CD106 antibody (#105718; BioLegend). After resuspension in PBS containing 2% FBS, the cells were sorted to collect the CD106-positive population using a BD FACS Aria II or SONY MA 900.

### Cell culture

C2C12 cells were obtained from ATCC and cultured in DMEM containing 10% FBS (GM) (Tsuchiya et al., 2018). MuSCs isolated from mice were grown on Matrigel-coated culture dishes (Corning) in growth medium (SC-GM; DMEM supplemented with 30% fetal bovine serum (Sigma-Aldrich), 1% chicken embryo extract (US Biological), 10 ng ml^−1^ basic fibroblast growth factor (#47079000; ORIENTAL YEAST Co., Ltd.), and 1% penicillin– streptomycin (FUJIFILM Wako Pure Chemical Corporation)) (Murakami et al., 2026). Cultures were maintained at 37°C in an incubator with 5% CO_2_.

### Real-time qPCR analysis

Total RNA was extracted from C2C12 cells or isolated MuSCs with ISOGEN II (#311-07361; Nippon Gene), and cDNA was synthesized using ReverTra Ace qPCR RT Master Mix (#FSQ-201; Toyobo). Total DNA was extracted using ISOGENOME (#314-08113; Nippon Gene). qPCR was conducted with PowerUp SYBR Green Master Mix (Thermo Fisher Scientific) on the StepOne system (#A25742; Thermo Fisher Scientific). Relative expression levels were calculated by the 2-ΔΔCt method. The primer sequences are provided in the Supplementary Table.

### Myotube formation

C2C12 cells grown to semiconfluence in 10% FBS/DMEM were transferred to differentiation medium (DM; DMEM containing 2% horse serum (Gibco)) for the indicated periods. For the 4-day incubation, the medium was replaced with fresh DM 2 days after the start of differentiation. MuSCs cultured in SC-GM for 6 days after isolation were transferred to differentiation medium (SC-DM; 5% horse serum and 1% penicillin–streptomycin) for a further 2 days (Murakami et al., 2026).

### RNA interference using siRNA

C2C12 cells and MuSCs were transfected using Lipofectamine RNAiMAX (#13778150; Thermo Fisher Scientific) with 10 μM siRNA (#SASI_Mm01_00186905, #SIC001, Sigma-Aldrich; #s80544, Thermo Fisher). C2C12 cells received siRNA in GM for 16 h and were then cultured in siRNA-containing DM for the indicated periods. MuSCs received 3-h siRNA treatments in SC-GM on days 4 and 5 after isolation, followed by culture for 2 days in siRNA-free SC-DM.

### Lentivirus production and transduction

Lentiviruses were produced using a published protocol with minor modifications (Joung et al., 2017). Briefly, recombinant lentiviruses carrying *egfp* or mouse *Mtch2* fused to *egfp* were generated by transfecting Lenti-X 293T cells with pLJM1-EGFP or pLJM1-MTCH2-EGFP, respectively. For infection, C2C12 cells were plated in a 12-well plate, and lentiviral solution was added to each well. The plate was then centrifuged at 1,000 × g and 33°C for 2 h (spinfection). Transduced cells were analyzed 2 days after infection (Murakami et al., 2026).

### Immunofluorescent analysis

Cells were fixed in 4% paraformaldehyde (PFA)/PBS, permeabilized with 0.5% Triton X-100/PBS, blocked with 1% BSA/PBS, and incubated with antibodies. To assess myotube formation, we stained cells with anti-MyHC antibody (#14-6503-82; dilution 1:200, eBioscience) and DAPI (Dojindo) and imaged them with an epifluorescence microscope (Axio-Observer. Z1, Zeiss) equipped with a 10× objective lens. DAPI-stained nuclei were counted in ImageJ to calculate the fusion index, defined as the percentage of nuclei in MyHC+ cells relative to the total number of nuclei in each microscopic field. To assess MuSC differentiation, we stained cells with anti-Myogenin antibody (#sc-576; dilution 1:500, Santa Cruz) and DAPI (Dojindo) and imaged them with an epifluorescence microscope (Axio-Observer. Z1, Zeiss) equipped with a 10× objective lens. DAPI-stained nuclei were counted in ImageJ, and the percentage of nuclei within multinucleated myosin heavy chain-positive myotubes was quantified in each microscopic field. This percentage was defined as the fusion index.

### Evaluation of cytotoxicity

C2C12 cells were incubated in 48-well plates for 48 h in 300 μL of culture medium with or without *Mtch2* siRNA. To assess the number of viable cells, we added 40 μL of CellTiter AQueous One (#G3582; PROMEGA) to each well. Following incubation at 37°C for 1 h, absorbance was measured at 490 nm with an infinite F200 PRO (Tecan).

### Evaluation of MuSC proliferation

Isolated MuSCs were incubated for 3 h in 24-well plates with GM containing 5 mM EdU. Incorporated EdU was detected by a click chemistry reaction following primary and secondary staining, using a Click-iT EdU Imaging Kit (#C10086; Thermo Fisher Scientific) according to the manufacturer’s instructions.

### RNA-seq analysis

After 2 days of culture under low-serum conditions, MuSC-derived myoblasts were harvested and pelleted by centrifugation. The pellets were snap-frozen in liquid nitrogen and sent to KOTAI Biotechnologies (Japan), where RNA extraction, library preparation, RNA sequencing, and subsequent bioinformatic analyses were performed.

### Observation of mitochondrial morphology

Mitochondria in C2C12 cells or MuSCs were visualized with Mitotracker Green FM (#M7514; Thermo Fisher Scientific). Cells were incubated with Mitotracker reagent (300 nM) in normal growth medium for 30 min at 37°C, then washed twice with PBS. During imaging, cells were maintained in serum-free DMEM containing HEPES (FUJIFILM Wako Pure Chemical Corporation). For Fig. 5A, mitochondria in MuSCs isolated from *Mtch2* cKO mice were visualized by immunofluorescence staining with anti-TOM20 antibody (#42406S; dilution 1:200, Cell Signaling Technology). Fluorescence images were acquired with a confocal laser-scanning microscope (LSM 800; Carl Zeiss) equipped with a 63× objective lens, a paired argon laser (488-nm line), and a bandpass filter (500-700 nm).

### Measurement of metabolic activity using an extracellular flux analyzer

The oxygen consumption rate (OCR) of C2C12 cells or isolated MuSCs was measured in 8-well plates using a Seahorse XFe Extracellular Flux Analyzer (Agilent). C2C12 cells and isolated MuSCs were seeded in Seahorse 8-well plates coated with poly-L-lysine and Matrigel, respectively. For the mitochondrial stress test (Fig. 3B and C, S4B and C), the medium was replaced before measurement with XF assay medium containing 5 mM or 25 mM glucose and 1 mM pyruvate. Cells were treated with 3 μM oligomycin, followed by 1 μM FCCP and a mixture of 0.5 μM rotenone and 0.5 μM antimycin. For the glycolysis stress test (Fig. 4A), the medium was replaced before measurement with XF assay medium containing 2 mM glutamate. Cells were treated sequentially with 10 mM glucose, 3 μM oligomycin, and 50 mM 2-deoxy-D-glucose (2-DG). For the palmitate oxidation stress test (Fig. 4B), cells were preincubated for 1 h at 37°C in XF assay medium containing 0.5 mM glucose, 1 mM glutamate, 0.5 mM L-carnitine, and 1% FBS. The medium was then replaced with XF assay medium containing 2 mM glucose, 0.5 mM L-carnitine, and 166 μM palmitate. Cells were subsequently treated with 60 μM etomoxir and a mixture of 0.5 μM rotenone and 0.5 μM antimycin at the indicated concentration. Oligomycin inhibits mitochondrial F_1_F_o_-ATPase, suppressing ATP synthesis-dependent OCR. FCCP is a chemical uncoupler that maximizes the electron transfer activity of mitochondrial complexes I–IV independently of ATP synthesis. Rotenone and antimycin A inhibit mitochondrial complexes I and III, respectively. The glucose analog 2-deoxy-D-glucose acts as a competitive inhibitor of hexokinase, suppressing glycolysis-dependent OCR. Etomoxir inhibits mitochondrial CPT1, a rate-limiting enzyme of mitochondrial β-oxidation.

### Histological analysis

Cardiotoxin experiments were conducted as previously described (Hirano et al., 2023). CTX (50 μl of 10 μM) was injected into the TA muscle of mice aged 8–15 weeks. Muscle samples were collected at the time points indicated in each figure and snap-frozen in isopentane cooled with liquid nitrogen. Transverse cryosections (thickness, 7 μm) were stained with hematoxylin and eosin as previously described. The cross-sectional area (CSA) in each muscle sample was determined from cryosections stained with anti-laminin antibody (#L9393; Sigma-Aldrich; 1:500). CSA and fluorescence intensity were measured in ImageJ for statistical analysis.

### Statistical analysis

Statistical analyses were conducted in Microsoft Excel and R version 3.6.3 (2020-02-29). Differences between mean values were assessed for statistical significance using an unpaired, two-sided t-test. Multiple comparisons were assessed using Tukey’s test followed by analysis of variance. Statistical significance was denoted by \**P* < 0.05, \*\**P* < 0.01, and \*\*\**P* < 0.001.

## Acknowledgements

We thank members of the Hara laboratory for their scientific contributions. For providing the mutant mouse line (*Mtch2*^tm1a(EUCOMM)Hmgu^), we thank the Welcome Trust Sanger Institute Mouse Genetics Project (Sanger MGP) and its funders, and the European Mouse Mutant Archive (www.emmanet.org) partner Medical Research Council (MRC) from which the mouse line was received. Funding and associated primary phenotypic information may be found at www.sanger.ac.uk/mouseportal. We also thank Dr. Shigeyoshi Itohara and the RIKEN BRC for providing the B6-Tg(CAG-FLPe)36 mice (RBRC01834). The authors are grateful to KOTAI Biotechnologies, Inc. for their technical assistance in bulk RNA sequencing and data analysis. This study was supported by the Grant-in-Aid for Scientific Research KAKENHI (22H03484, 25K02994); Grant-in-Aid for Transformative Research Areas B (23H03854, 23H03856); Intramural Research Grant (5-6, 8-13) for Neurological and Psychiatric Disorder of NCNP; grants from Takeda Science Foundation, ONO Medical Research Foundation, Uehara memorial foundation, Chugai Foundation for Innovative Drug Discovery Science (SRG2022), and the Asahi Glass Foundation (to Y.H.); and the Grant-in-Aid for Scientific Research KAKENHI (22K20636, 23K14184), UBE Foundation, ONO Medical Research Foundation (to A. Murakami.).

## Author contributions

Conceptualization: Y.H., K.N., and M.U.; Methodology: K.H., M. Tsuchiya., Y.O.; Investigation: A. Murakami., Y.A., A. Miyazawa., T. Masuda and M.S.; Supervision: A. Murakami., Y.H.; Writing—original draft: A. Murakami.; Writing—review and editing: A. Murakami., Y.H.

## Data and material availability

All data needed to evaluate the conclusions in the paper are present in the paper.

