## Supplementary figures 1-8, Supplementary Table for "MTCH2 drives metabolic remodeling in muscle satellite cells via mitochondrial fatty acid utilization"

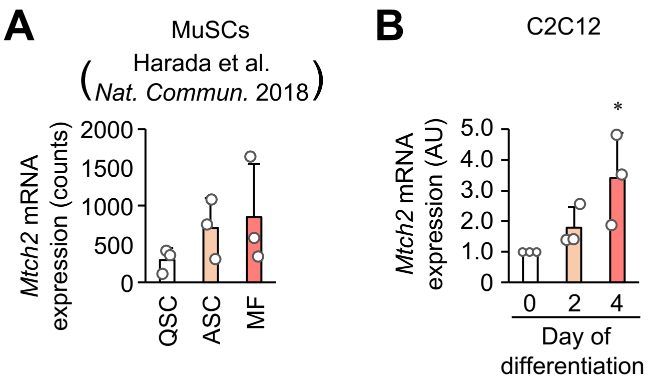


Murakami, A et al. Supplementary Figure. 1

**Supplementary Figure 1.** ***Mtch2*** **expression during myogenesis.**

(A) Changes in *Mtch2* mRNA expression in MuSCs following the induction of skeletal muscle injury. Analyses were conducted *in silico* with previously reported RNA-seq data. QSC, quiescent satellite cell; ASC, activated satellite cell; MF, myofiber. (B) Changes in *Mtch2* mRNA expression in C2C12 cells during culture in low-serum medium for the indicated periods. Data are presented as mean + SD. n = 3. *$P < 0.05$.

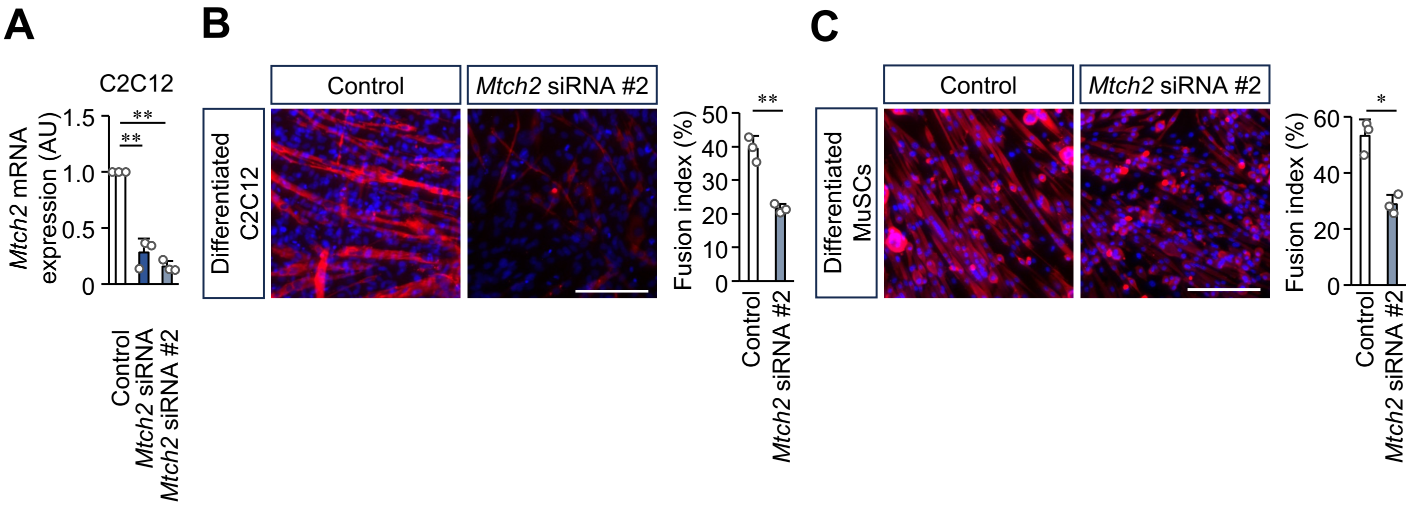


Murakami, A et al. Supplementary Figure. 2

**Supplementary Figure 2. Myogenic differentiation in myoblasts treated with independent *Mtch2* siRNAs.**

(A) *Mtch2* mRNA expression in C2C12 cells following treatment with two independent *Mtch2* siRNAs. (B) Myotube formation in C2C12 cells treated with *Mtch2* siRNA #2 and cultured for 4 days in low-serum medium. (C) Myotube formation in myoblasts derived from isolated MuSCs following treatment with Mtch2 siRNA #2 and culture for 2 days in low-serum medium. Myotube formation was assessed using the fusion index. Red, myosin heavy chain; blue, nuclei. Scale bars, 100 μm. Data are presented as mean + SD. n = 3. *$P < 0.05$, **$P < 0.01$, ***$P < 0.001$.

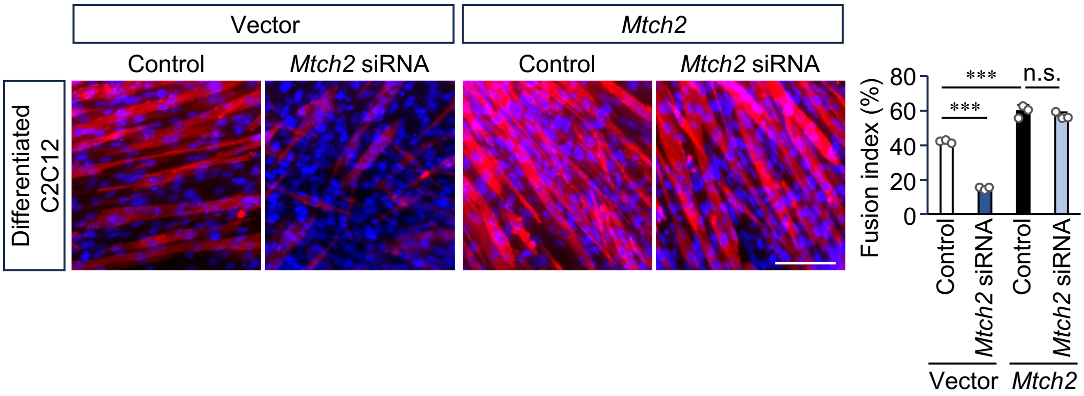


Murakami, A et al. Supplementary Figure. 3

**Supplementary Figure 3. Myotube formation in *Mtch2*-overexpressing myoblasts.**

*Mtch2* was overexpressed by lentiviral transduction in C2C12 cells maintained under growth conditions. To induce myotube formation, cells were cultured for 4 days in low-serum medium in the presence of *Mtch2* siRNA. Myotube formation was assessed using the fusion index. Red, myosin heavy chain; blue, nuclei. Scale bar, 100 μm. Data are presented as mean + SD. n = 3. ***$P < 0.001$; n.s., not significant.

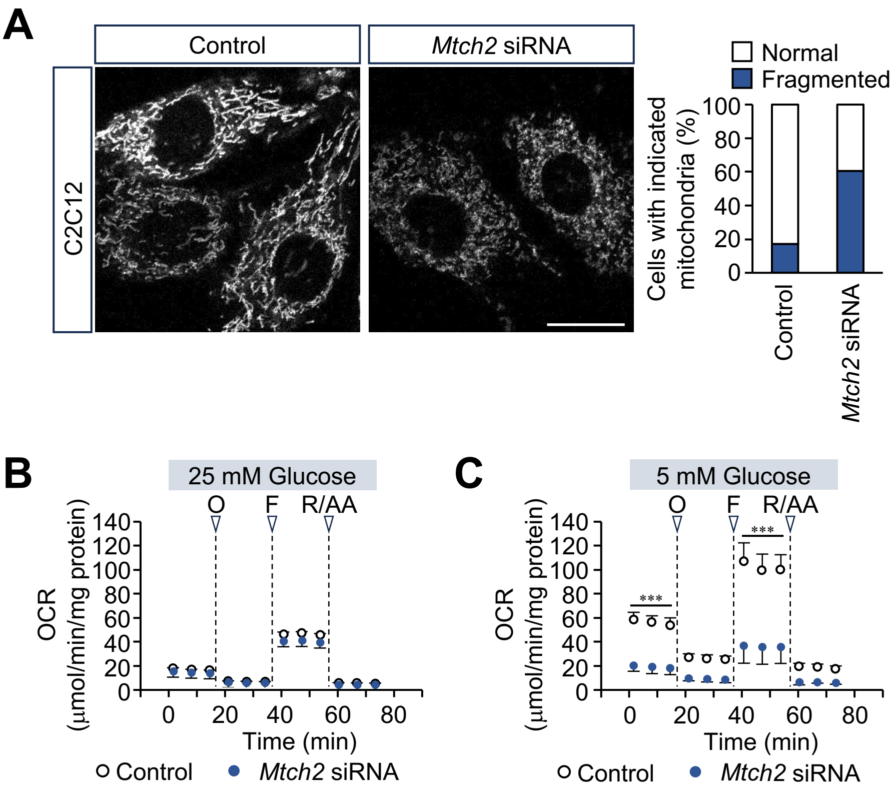


Murakami, A et al. Supplementary Figure. 4

**Supplementary Figure 4. Mitochondrial morphology and respiration in *Mtch2*-depleted C2C12 cells.**

(A) Mitochondrial morphology. Following MitoTracker® staining, mitochondria were classified as fused/elongated (Normal) or fragmented (Fragmented). Control, n = 189; *Mtch2* siRNA, n = 175. Scale bar, 20 µm. (B, C) Oxygen consumption rate (OCR) measured in assay medium containing 25 mM glucose (B) or 5 mM glucose (C). O, oligomycin; F, FCCP (carbonyl cyanide 4-(trifluoromethoxy)phenylhydrazone); R/AA, rotenone/antimycin A. *Mtch2* depletion was achieved using siRNA. Data are presented as mean + SD. n = 3. ***$P < 0.001$.

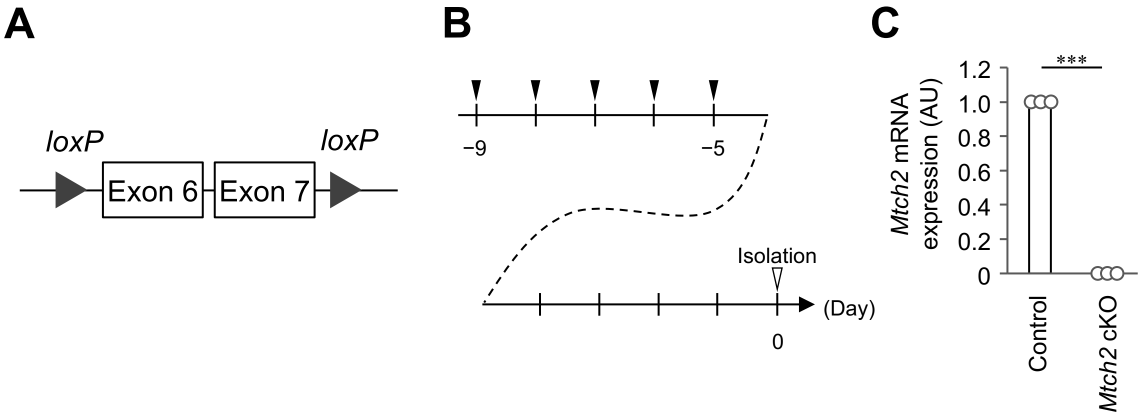


Murakami, A et al. Supplementary Figure. 5

**Supplementary Figure 5. MuSC-specific *Mtch2* conditional knockout (*Mtch2* cKO) mice.**

(A) Schematic of the *Mtch2* locus in *Mtch2*$^{flox/flox}$ mice. (B) Experimental scheme showing MuSC-specific *Mtch2* knockout. Black arrows, intraperitoneal tamoxifen administration. (C) Expression of *Mtch2* mRNA measured 6 days after isolation of MuSCs from *Mtch2* cKO mice. Data are presented as mean + SD. N = 3 mice. ****P* < 0.001.

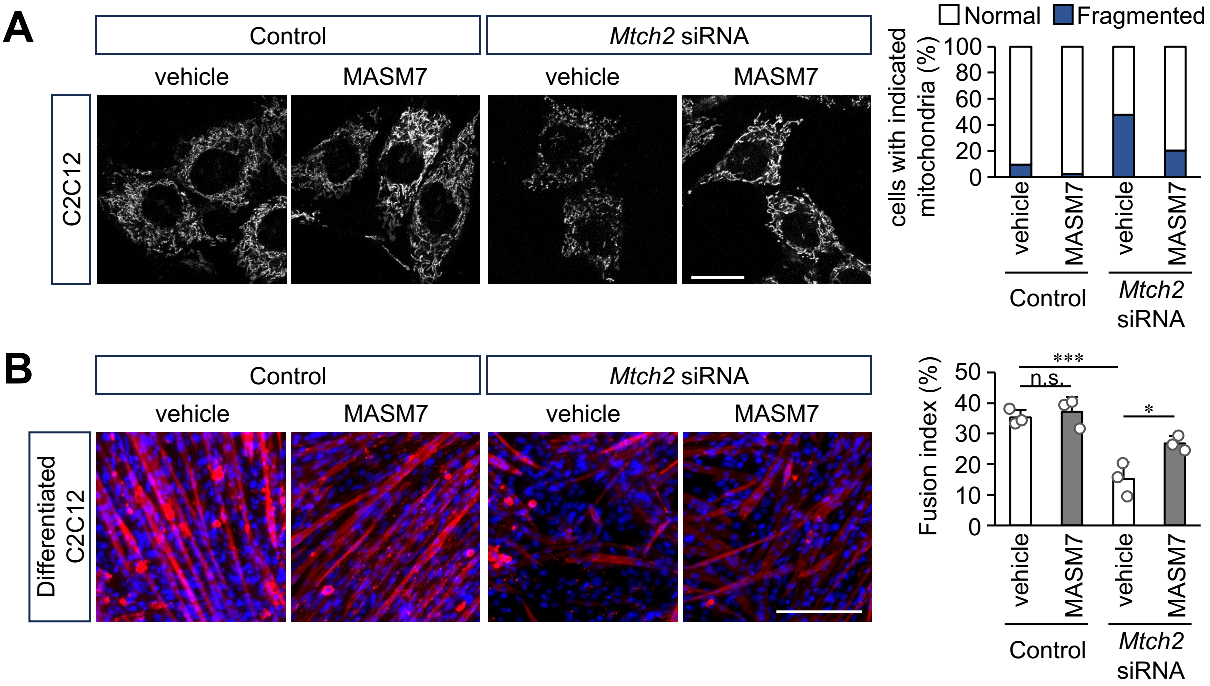


Murakami, A et al. Supplementary Figure. 6

**Supplementary Figure 6. Dependence of myotube formation on the regulation of mitochondrial morphology.** (A) Effect of the mitochondrial fusion activator MASM7 (5 μM) on mitochondrial fragmentation induced by *Mtch2* siRNA in C2C12 cells. Mitochondria were stained with MitoTracker® and classified as fused/elongated (Normal) or fragmented (Fragmented). Control/MASM7(−), n = 154; Control/MASM7(+), n = 154; *Mtch2* siRNA/MASM7(−), n = 180; *Mtch2* siRNA/MASM7(+), n = 190. Scale bar, 20 μm. (B) Effect of MASM7 (5 μM) on the suppression of myotube formation induced by *Mtch2* siRNA in C2C12 cells. Myotube formation was assessed using the fusion index. Red, myosin heavy chain; blue, nuclei. Scale bar, 100 μm. Data are presented as mean + SD. n = 3. *$P$ < 0.05, ***$P$ < 0.001; n.s., not significant.

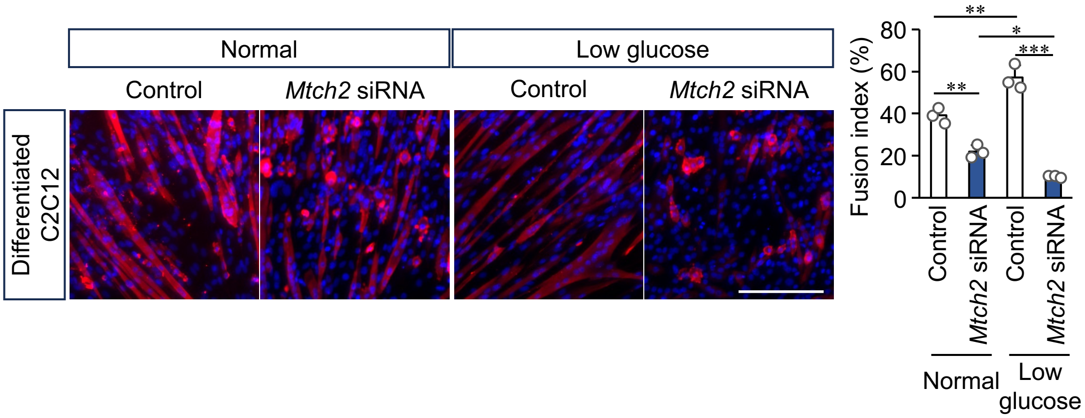


Murakami, A et al. Supplementary Figure. 7

**Supplementary Figure 7. Myotube formation under low-glucose conditions.**

To induce myotube formation, C2C12 cells were cultured for 4 days in low-serum medium containing 25 mM glucose (Normal) or 5 mM glucose (Low glucose) in the presence of *Mtch2* siRNA. Myotube formation was assessed using the fusion index. Red, myosin heavy chain; blue, nuclei. Scale bar, 100 μm. Data are presented as mean + SD. n = 3. $*P < 0.05$, $**P < 0.01$, $***P < 0.001$.

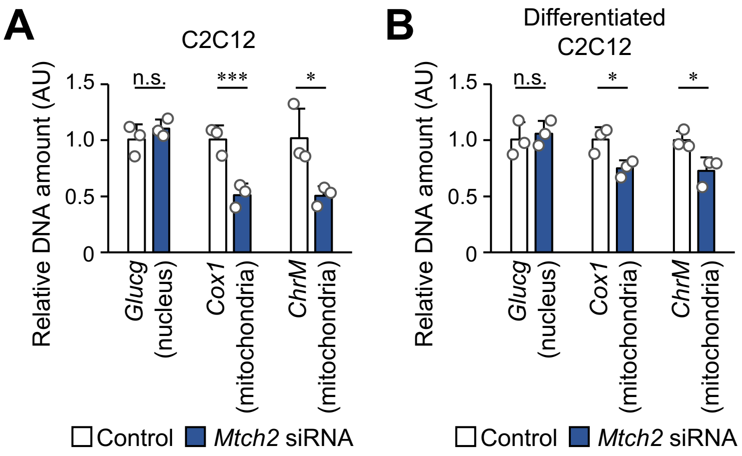


Murakami, A et al. Supplementary Figure. 8

**Supplementary Figure 8. Evaluation of mitochondrial mass in *Mtch2*-depleted myoblasts.**

Nuclear- and mitochondrial-localized genomic DNA amounts were measured by quantitative genomic PCR. C2C12 cells were cultured under normal growth conditions (A) or for 4 days under low-serum conditions (B) before analysis. *Mtch2* depletion was achieved using siRNA. Data are presented as mean + SD. n = 3. *$P < 0.05$, ***$P < 0.001$; n.s., not significant.

**Supplementary Table**

Primers used for quantitative real-time PCR in this study.

| | |
|---|---|
| *18S_F* | TTCTGGCCAACGGTCTAGACAAC |
| *18S_R* | CCAGTGGTCTTGGTGTGCTGA |
| *Mtch2_F* | CCACGTGATCACTCTGAGGT |
| *Mtch2_R* | ACCTAGGAGGCGAGGAATGA |
| *Myog_F* | CGCCATCCAGTACATTGAGCG |
| *Myog_R* | AGGAGGCGCTGTGGGAGTT |
| *Myh4_F* | GCAGGACTTGGTGGACAAAC |
| *Myh4_R* | ACTTGGCCAGGTTGACATTG |
| *Mck_F* | ACCTCCACAGCACAGACAGA |
| *Mck_R* | CAGCTTGAACTTGTTGTGGG |
| *Mymk_F* | GACAGTGAGCATCGCTACCA |
| *Mymk_R* | GCACAGCACAGACAAACCAG |
| *Glucg_F* | CAGGGCCATCTCAGAACC |
| *Glucg_R* | GCTATTGGAAAGCCTCTTGC |
| *Cox1_F* | AGCCCACTTCGCCATCATAT |
| *Cox1_R* | GCGTCGTGGTATTCCTGAAAG |
| *ChrM_F* | AATCAACTCGTCTATGTGGCAAAA |
| *ChrM_R* | CCAGCTATCACCAAGCTCGTT |
| *Uqcr11_F* | TCTGCACATGCGTAGTGCTC |
| *Uqcr11_R* | CGGCTGTGGGAATCCAGTTT |
| *Uqcrq_F* | AGAGCTCTGGAGGGCCG |
| *Uqcrq_R* | TGGGGATGCCTTTGCTGAAA |
| *Cox8a_F* | CAGCTCGGGGTCCTGGATA |
| *Cox8a_R* | ACTCCCGCTTCTTGTAGCTC |
| *Ndufb8_F* | CCTACCAGCCTGTGGAATAGT |
| *Ndufb8_R* | GCAGGAAAACAGGAATGCAGG |
| *Chchd10_F* | GTGCAAATACAATCACGGTCTGA |
| *Chchd10_R* | CTCCCAGTCACATGGAACCT |
| *Atg101_F* | TACTGCGTGTTCGCGTTAGG |
| *Atg101_R* | ACAGGATACTTCACCCCACCT |

| | |
|---|---|
| *Enho_F* | ATCTCCCAAGGGGCTCTCAT |
| *Enho_F* | CGACGTCAGCAGATCGAGAA |
| *Galk1a_F* | GCACACGGACTACAACCAGG |
| *Galk1a_R* | GAAACAAGCCCATCTGTCCG |
| *Higd1a_F* | GAAGGGACCCCGCGTCT |
| *Higd1a_R* | TTCCAATGGGGACAAACGGT |
| *Pgls_F* | CTTCGACCACGCCGAGAG |
| *Pgls_R* | TGGTAAGGACCTGGCTGTCT |
